# RNA interactions drive structural and functional diversification of α-Synuclein fibrils

**DOI:** 10.1101/2025.11.30.691421

**Authors:** Jakob Rupert, Francesco di Palma, Alessio Colantoni, Antonia Intze, Maria Eleonora Temperini, Valeria Giliberti, Elsa Zacco, Gian Gaetano Tartaglia

## Abstract

Nucleic acids, particularly RNA, exert multiple roles on protein aggregation, yet their mechanistic involvement remains poorly understood. Building on prior work demonstrating RNA-driven acceleration of protein aggregation, we now demonstrate that RNA binding markedly reshapes the structural and functional properties of alpha-synuclein (aS) fibrils, a protein implicated in Parkinson’s disease (PD). Although monomeric aS exhibits negligible RNA-binding properties, aggregation markedly increases its RNA-binding capacity. Our kinetic analyses reveal that RNA incorporation during initial stages of aggregation alters fibril elongation dynamics and modulates surface-mediated secondary nucleation in a seed concentration-dependent manner. Spectroscopic and microscopic characterization (FTIR, micro-FTIR) demonstrate that RNA promotes the formation of structurally distinct, more compact aS fibrils with heterogeneous morphology. Exploiting the recently discovered catalytic hydrolase activity of aS fibrils, we show that direct RNA binding inhibits enzymatic function, consistent with RNA-induced conformational remodeling that limits the exposure of critical residues. Additionally, RNA binding to fibril surfaces impairs proteolytic degradation by proteinase K, suggesting RNA stabilizes fibrils against clearance mechanisms. Moreover, RNA sequencing of fibril-bound nucleic acids uncovers a preference towards uridine-rich motifs that computational docking locates within hydrophilic cavities of disease-associated fibril structures. Collectively, we identify RNA as an active co-factor that remodels structure, propagation kinetics, stability, and function of aS fibrils. We offer a mechanistic link between RNA interaction and amyloid polymorphism relevant to synucleinopathies and, in broader contexts, to amyloid-associated diseases.

**Significance statement:** Our work represents the first systematic evidence that RNA directly interacts with protein fibrils and modulates key functional properties such as aggregation kinetics, proteolytic resistance and catalytic activity. We demonstrate that RNA co-aggregates with alpha-synuclein inducing structural conformations that depend on RNA for efficient assembly. Given the resemblance of these RNA-associated fibrils to those found in multiple system atrophy, our findings suggest that the final fibril structures in the patient brain are influenced by multiple cofactors and may differ substantially from recombinant fibrils formed *in vitro*, even when seeded with patient-derived material.

## Introduction

Alpha-synuclein (aS) is a presynaptic, intrinsically disordered protein implicated in several neurodegenerative disorders, including Parkinson’s disease (PD) and multiple system atrophy (MSA) (1). These conditions are marked by the accumulation of aS into amyloid fibrils, which assemble into intracellular inclusions such as Lewy bodies and glial cytoplasmic inclusions. Traditionally, the toxic gain-of-function in aS pathology has been attributed to fibrillar aggregation. However, recent studies challenge this view. Mahul-Mellier *et al.* demonstrated that fibril presence alone is insufficient to recapitulate disease features, proposing instead that Lewy body formation is a multistep process involving the recruitment of organelles, lipids, and nucleic acids into a structured inclusion—this being the true pathological hallmark (2).

This perspective raises critical questions about the cellular factors modulating aS aggregation. Among these, nucleic acids have emerged as potent modulators of protein assembly. Increasing evidence shows that RNA can influence the kinetics and structure of aggregates formed by proteins such as tau, FUS, and TDP-43 (3, 4). In our previous work, we demonstrated that aggregation confers an acquired RNA-binding ability to aS—a property absent in its monomeric state (5). We showed that removal of the acidic C-terminal region enhances both aggregation and RNA sequestration, due to favorable electrostatics and increased exposure of the positively charged N-terminus, particularly around residue H50. Although RNA-mediated aggregation of aS is clearly dependent on RNA concentration, the contribution of sequence specificity remains unexplored. It is plausible that certain RNA motifs, such as G-quadruplexes (G4), might play a role in the early stages of aggregation (6).

Previous computational analysis of diverse amyloid-forming proteins revealed a common architecture: hydrophobic, aggregation-prone cores buried within fibrils, flanked by disordered, solvent-exposed regions capable of interacting with nucleic acids (5). These external domains can contain RNA-binding motifs or electrostatic patches that promote interactions with RNA backbone. For aS, the N-terminal region shows the highest RNA-binding propensity, consistent with cryo-electron microscopy (cryo-) studies of MSA-derived fibrils, identifying unassigned electron densities within positively charged cavities formed by residues K43, K45, and H50 (7).

Building on these observations, this study investigates the functional and structural consequences of RNA binding to aS aggregates. We examined how RNA modulates aggregation kinetics, alters fibril structure, enhances protease resistance, and suppresses the hydrolase activity recently attributed to mature aS fibrils. By estimating RNA-binding affinities and proposing an inhibition model, we reveal a nuanced role for RNA as a cofactor that interacts with fibrils post-aggregation and alters their templated growth. Considering our results alongside previous reports on RNA colocalization with protein aggregates in disease tissue (e.g., tau, hnRNPDL-2), we propose that RNA contributes actively to the structural and functional diversity of aS aggregates (8). Understanding the interplay between RNA and aS fibrils may help explain the emergence of disease-specific polymorphs and the selective vulnerability observed in synucleinopathies. It also opens new avenues for RNA-targeted therapeutic interventions in disorders marked by pathological protein aggregation.

## Results

### RNA alters aS aggregate propagation via seed-specific mechanisms

Primary nucleation is the main mechanism driving aS aggregation under near-physiological conditions (9), and previous work has shown that RNA significantly accelerates this process in a concentration-dependent manner (5). To enhance the reproducibility of our results and control changes in lag time, we initially performed aggregation assays using glass beads and continuous shaking. However, such conditions are known to promote fibril fragmentation and artificially elevate both primary and secondary nucleation rates, as observed for Aβ42 (10). To reduce these shear-induced effects, we subsequently conducted *in vitro* seeded aggregation assays under quiescent conditions, employing seed concentrations high enough to saturate primary nucleation. In this regime, fibril growth is primarily driven by seed elongation and surface-mediated secondary nucleation, consistent with recent observations under physiological conditions (11).

We then compared the seeding efficiency of aggregates formed in the absence or presence of RNA, referred to as protein^only^ and protein^RNA^ seeds, respectively. We aimed to determine how RNA incorporated during initial aggregation affects subsequent fibril growth in the presence of additional protein monomers and RNA. All conditions exhibited exponential fluorescence increases without a lag phase (**Fig. 1A**), consistent with saturated primary nucleation and elongation-driven growth. Since elongation is a first-order process dependent on fibril ends and monomer concentration, we extracted kinetic parameters by fitting early-phase data (*t* < 1 h) with a linear model and compared slopes across conditions (**Supp. Fig. 1A**). This enabled us to directly assess elongation efficiencies of the different seed populations. Our results reveal that protein^only^ seeds consistently result in higher elongation rates than protein^RNA^ seeds, (**Fig. 1B**), suggesting RNA incorporation during fibril formation reduces the seeds’ ability to promote monomer addition. This trend is supported by lower endpoint fluorescence values for protein^RNA^ seeds (**Supp. Fig. 1B**), indicating reduced total fibril mass. These findings suggest that RNA structurally alters aS aggregates, impairing their seeding capacity.

**Figure 1.**
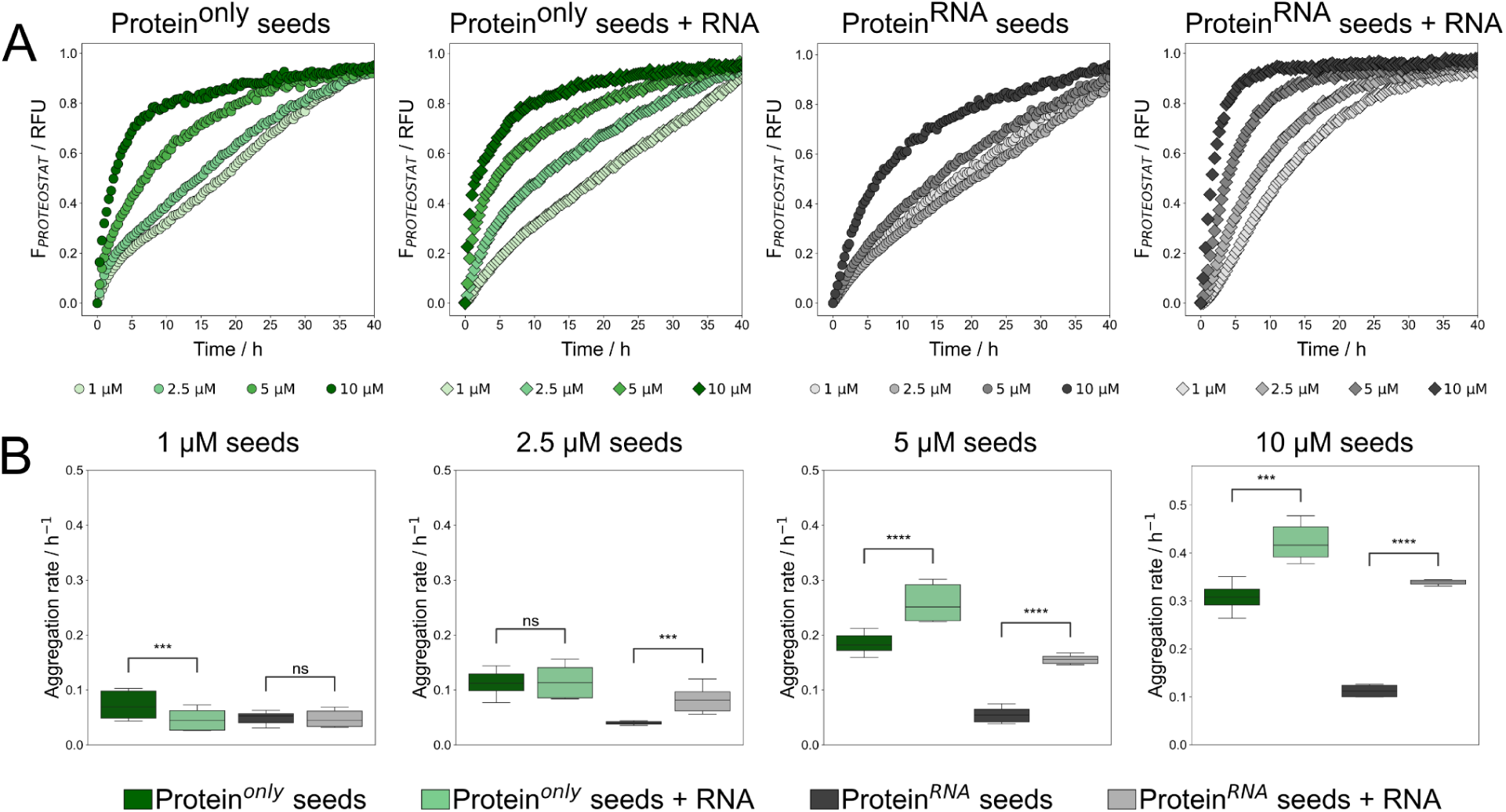
Seeded aggregation assays performed under quiescent conditions using aS seeds generated in the absence (protein^only^ seeds) or presence of RNA (protein^RNA^ seeds). (*A*) Proteostat™ fluorescence signal of 50 μM aS subjected to seeded aggregation with protein^only^ and protein^RNA^ seeds at varying concentrations, in the presence or absence of RNA. All traces exhibit an exponential profile with no evident lag phase, indicating saturation of primary nucleation processes. Protein^only^ seeds show higher seeding efficiency compared to protein^RNA^ seeds across all conditions. Notably, the addition of RNA markedly accelerates aggregation in the case of protein^RNA^ seeds. At lower seed concentrations, the traces exhibit increased complexity, suggestive of overlapping aggregation mechanisms. The initial phase of the reaction shows a linear increase in fluorescence, consistent with elongation-driven fibril growth. (*B*) Analysis of the initial elongation rates reveals that at higher seed concentrations (5 μM and 10 μM), RNA significantly enhances the rate of fibril growth (*p* < 0.0001, *N* = 8, two-way Student’s *t*-test). Conversely, at 1 μM seed concentration, protein^only^ seeds exhibit significantly faster elongation in the absence of RNA (*p* < 10⁻⁵, *N* = 8), while no significant difference is observed for protein^RNA^ seeds under the same condition.

To further analyze kinetics, we fit late-stage data (t > 10 h) with a Hill’s function (see **Materials and Methods**). At lower seed concentrations, protein^only^ seeds in the absence of RNA showed a higher Hill’s coefficient (h = 1.77 ± 0.67) compared to those with RNA (h = 1.12 ± 0.31), indicating more prominent secondary processes (**Supp. Fig 2A)**. Thus, fibril growth without primary nucleation is driven by elongation and secondary nucleation, resulting in a higher apparent initial aggregation rate. Importantly, RNA appears to have a dual role: accelerating elongation while reducing or blocking surface-mediated secondary nucleation (**Fig. 2A**). A different trend is seen with protein^RNA^ seeds: elongation rates show minimal change at lower concentrations of RNA but increase at 10 μM (**Fig. 2B**). Here, RNA significantly boosts elongation, with rates scaling linearly with seed concentration. This suggests that RNA is required for efficient growth of protein^RNA^ fibrils and supports the idea that these fibrils are structurally distinct from protein^only^ fibrils. Additionally, we showed that aS sequesters RNA during aggregation (5). Consistent with this, we found lower soluble RNA levels after aggregation with protein^RNA^ seeds than with protein^only^ seeds (p < 0.001, N = 8, **Supp. Fig. 2B**). This reinforces RNA’s role in promoting RNA-templated fibril propagation. Co-aggregation with RNA appears to enhance fibril RNA-binding capacity, promoting further RNA incorporation. In contrast, protein^only^ seeds sequester less RNA, suggesting their architecture is less compatible with nucleic acid binding.

**Figure 2.**
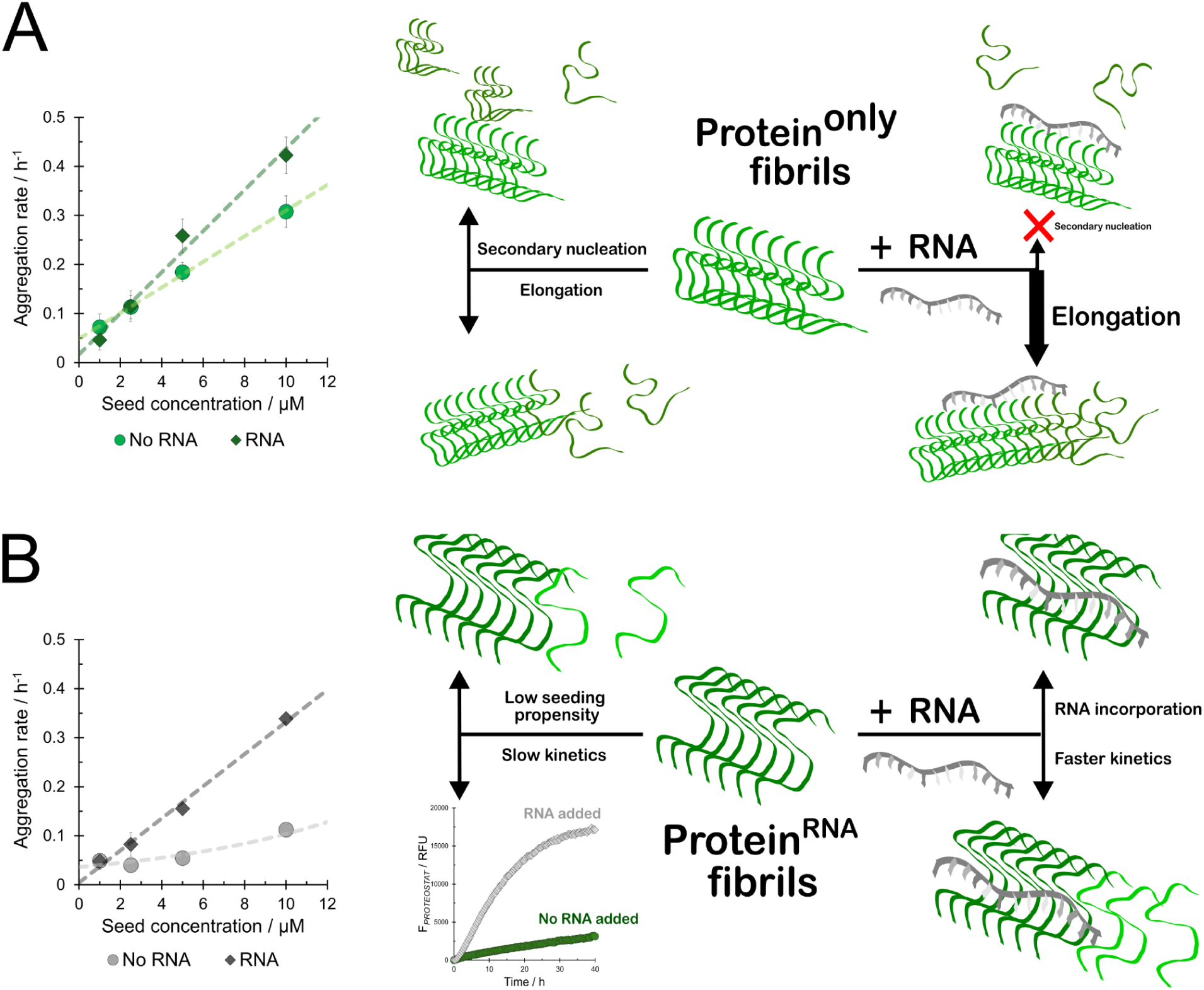
RNA affects the seeded growth of aS fibrils via different mechanisms. (*A*) Plotting the rates against seed concentration indeed reveals different trends for protein^only^ seeds, with RNA impeding the elongation rate at lower and accelerating it at higher seed concentrations. (*B*) Protein^RNA^ seeds on the other hand exhibit very poor seeding capacity at lower seed concentrations, which increases dramatically when RNA is present in solution.

Together, these findings highlight RNA as an active cofactor that reshapes the structure, function, and propagation of aS aggregates.

### RNA changes the structural properties of aS aggregates

To assess the impact of RNA on the structure of protein^RNA^ seeded fibrils compared to protein^only^ seeded fibrils, we used Fourier-Transform Infrared (FTIR) spectroscopy (12). Spectra recorded from mm^2^-sized areas of sample deposited on CaF_2_ substrate revealed a strong β-sheet peak around 1630 cm^-1^ (here referred to as cross-β peak) in the amide I region (1600 – 1700 cm^-1^) (**Fig. 3A, top**), characteristic of amyloid cross-β structures (13). In protein^RNA^ fibrils, this peak was shifted to lower central frequencies (ω_cross-β_), as confirmed by second derivative analysis (**Fig. 3A, bottom**). Such a shift suggests a more compact fibrillar structure that can result from stronger hydrogen bonding (14).

**Figure 3.**
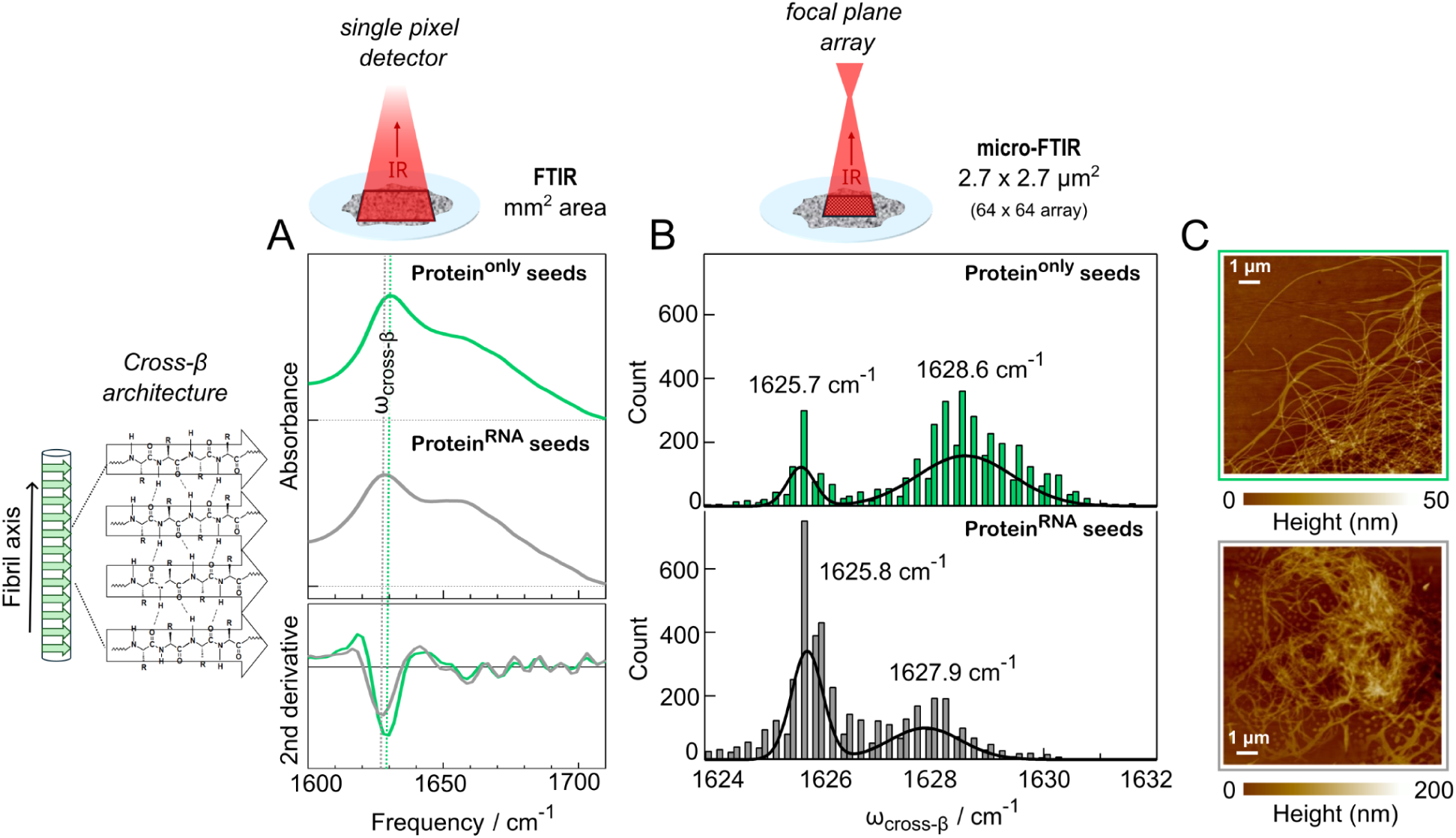
Characterisation of the structural properties of aS fibrils amplified from protein^only^ (green) and protein^RNA^ (grey) seeds. (*A*) Analysis of the shape and position of the cross-β peak for the two species shows a slight shift of the central peak frequency (ω_cross-β_) towards lower wavenumbers. (*B*) A similar shift towards lower wavenumbers can be observed from the distribution of the central frequencies amongst the sample micro-FTIR spectra, recorded with FPA detector. For protein^RNA^ seeds, the resulting fibrils have a much higher population of the conformation with the ω_cross-β_ at 1625.8 cm^-1^ compared to protein^only^ fibrils, indicating a reduced polymorphism that favors the more rigid fibril population . *(C)* Representative AFM maps of the protein^only^ and protein^RNA^ fibrils.

Because spectra from larger sample areas may average over diverse fibril conformations and cannot be exhaustively representative of the degree of polymorphism and possible effect of RNA on it, we repeated the experiment using FTIR microspectroscopy (micro-FTIR) with a focal plane array (FPA) detector. This setup allowed spectral acquisition from 2.7 x 2.7 μm^2^ regions across a 64 x 64 pixel array, enabling statistical analysis of ω_cross-β_ frequencies at considerably smaller areas (**Fig. 3B**). Both protein^only^ and protein^RNA^ fibrils displayed a bimodal ω_cross-β_ distribution, indicating two main distinct polymorphs. The more compact polymorph (ω_cross-β_ ≈ 1625.8 cm^-1^) was significantly more prevalent in protein^RNA^ fibrils, while the protein^only^ fibrils show a more heterogeneous distribution.

To confirm these differences, we performed atomic force microscopy (AFM) imaging (**Fig. 3C**). Protein^only^ fibrils appeared relatively uniform and slightly curved, while protein^RNA^ fibrils were shorter, more heterogeneous, and tended to form clumps. These findings demonstrate that RNA alters the structural properties of aS amyloid fibrils, favoring the formation of more compact polymorphs with distinct supramolecular features.

### RNA binding to aS fibril inhibits proteinase K digestion

We then assessed whether the increased structural compactness of RNA-induced aS fibrils could result in decreased proteolytic degradation, a key mechanism in cellular proteostasis and aggregate clearance (15). Since degradation involves direct interaction between the enzyme and fibril surface, we were also curious if incubating the fibrils with RNA could potentially block proteinase K (pK) cleavage.

Fibrils grown under quiescent conditions from protein^only^ seeds were incubated with pK and increasing RNA concentrations and analyzed by sodium dodecyl sulfate–polyacrylamide gel electrophoresis (SDS-PAGE). Higher RNA levels appeared to reduce proteolytic digestion (**Supp. Fig. 3A**). However, this method lacked quantitative resolution. To address this, we added 0.5 μg/mL pK directly to aggregation assay wells at the end of the reaction and monitored the fluorescence of the aggregate intercalator Proteostat^TM^, assuming successful digestion would lead to a fluorescence decrease. As in the gel analysis, increasing RNA concentrations slowed the fluorescence decay (**Fig. 4A**). Curve fittings revealed a decrease in initial reaction velocity with increasing RNA, following an exponential trend (**Supp. Fig. 3B**). In order to verify whether this reduction in activity was due to RNA binding the enzyme directly, we repeated the experiment using a standard enzymatic substrate pNPA with no fibrils present. Increasing concentrations of RNA did not inhibit pK activity in the absence of aS fibrils (**Supp. Fig. 3C**). This result supports the claim that RNA inhibits fibril degradation by binding to their surface and impeding the enzyme’s access to them.

**Figure 4.**
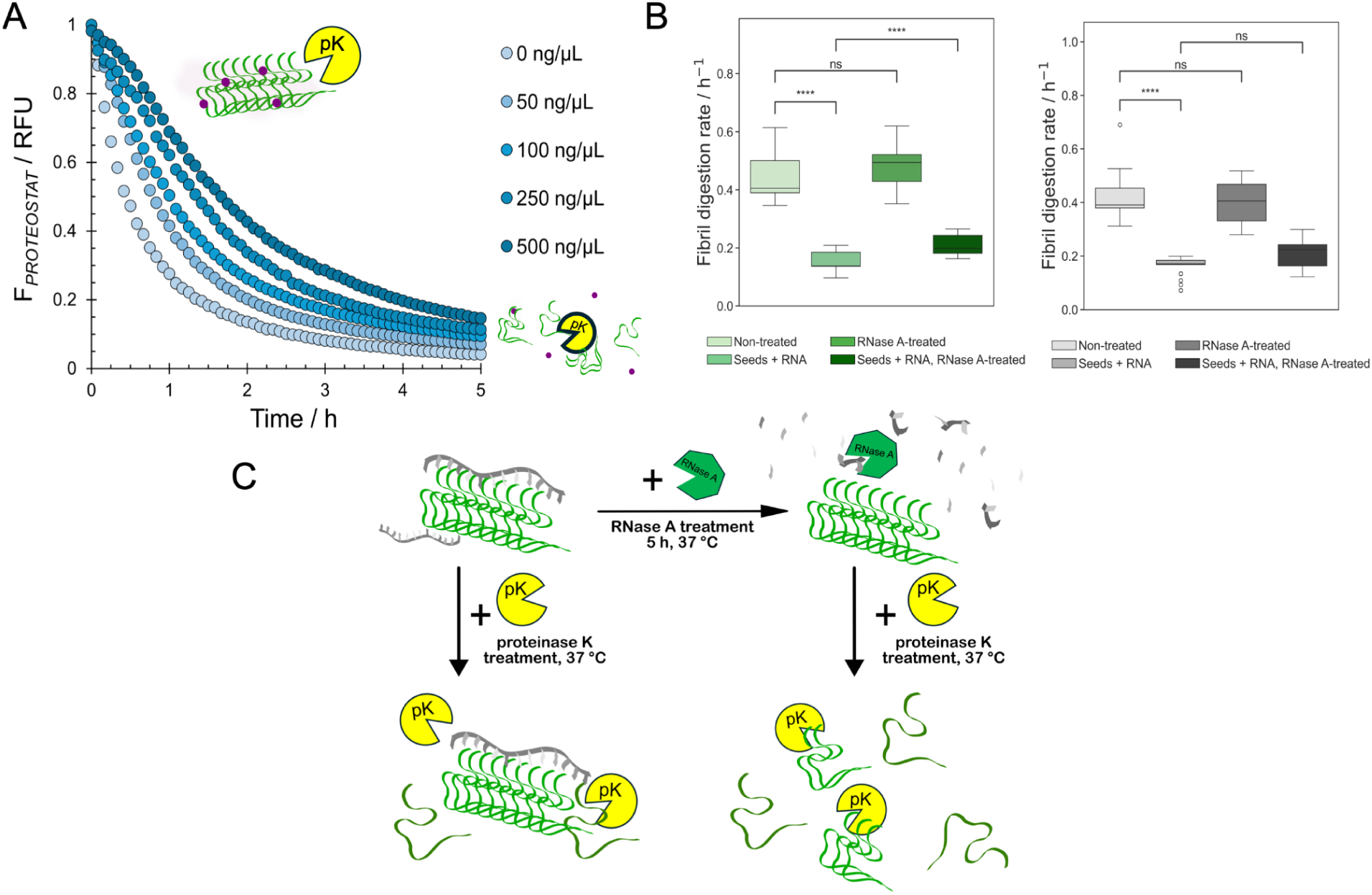
RNA impedes the efficient degradation of aS fibrils with proteinase. **K.** (*A*) Tracing the real-time Proteostat^TM^ fluorescence allowed for the quantification of the degradation kinetics at 0, 50, 100, 250, and 500 ng/μL RNA, indicating that incremental additions of RNA slow down the fluorescence decay, diminishing the reaction rate. (*B*) Comparing the reaction rates confirms the observation that there is no significant difference between reaction rates in the absence of RNA for both protein^only^ and protein^RNA^ fibrils, independent of RNase A treatment. RNA significantly decreases the degradation rate in both cases (p < 10^-10^, N = 30 (protein^only^ seeds), N = 24 (protein^RNA^ seeds), two-way Student t-test), while RNase A treatment significantly increases the rate of degradation for protein^only^ fibrils (p < 10^-5^, N = 30, two-way Student t-test). (*C*) These results confirm that the presence of RNA impedes the degradation of aS fibrils by pK and RNA degradation by RNase A ameliorates this effect, likely resulting in direct RNA binding and shielding of aggregated aS.

We hypothesized that if RNA binding impairs fibril digestion, removing RNA should restore proteolytic efficiency. To test this, we grew the fibrils from both protein^only^ and protein^RNA^ seeds in the presence or absence of RNA. RNase A or buffer was added, followed by pK, and fluorescence was monitored. RNase A or buffer alone had no effect on fluorescence signal (**Supp. Fig. 3D**). In fibrils grown without RNA, there was no difference in degradation rates between RNase A-treated and untreated samples (**Fig. 4B**). However, for protein^only^ fibrils grown with RNA, RNase A treatment led to significantly faster degradation (**Fig. 4B**), confirming that surface-bound RNA impedes proteolysis. In contrast, protein^RNA^ fibrils grown with RNA showed no significant difference in degradation rate following RNase A treatment (p = 0.06, N = 11). Notably, these fibrils exhibited a rapid initial fluorescence decrease that abruptly slowed. This suggests that in protein^only^ fibrils RNA remains surface-bound and directly blocks pK access, while in protein^RNA^ fibrils, RNA may be sequestered within the fibril core. Initial digestion might expose the sequestered RNA, subsequently inhibiting further degradation.

These findings indicate that RNA binding to fibril surfaces can impair proteolytic clearance (**Fig. 4C**). This mechanism may play a role in modulating amyloid stability *in vivo* and highlights the importance of cofactors in regulating amyloid proteostasis.

### RNA interacts directly with aS fibrils and inhibits their catalytic activity

With evidence that RNA inhibits pK digestion of aS fibrils in a concentration-dependent manner, we wanted to verify whether it was in fact the direct RNA-binding to fibril surface that was responsible. To demonstrate the ability of aS amyloid fibrils to interact directly with RNA, we leveraged a recent discovery that aS fibrils harbor catalytic hydrolase activity (16). The activity is linked to the only histidine in aS, H50, located in the region predicted to have high RNA-binding propensity (**Supp. Fig. 4**). In a form of aS aggregate (i.e., the “rod 1a” polymorph (17)), this region forms a pocket around H50 that closely resembles a similar RNA-binding pocket in RNA-induced tau fibrils (**Fig. 5A**)(4), as predicted by the *cat*RAPID approach (18, 19).

**Figure 5.**
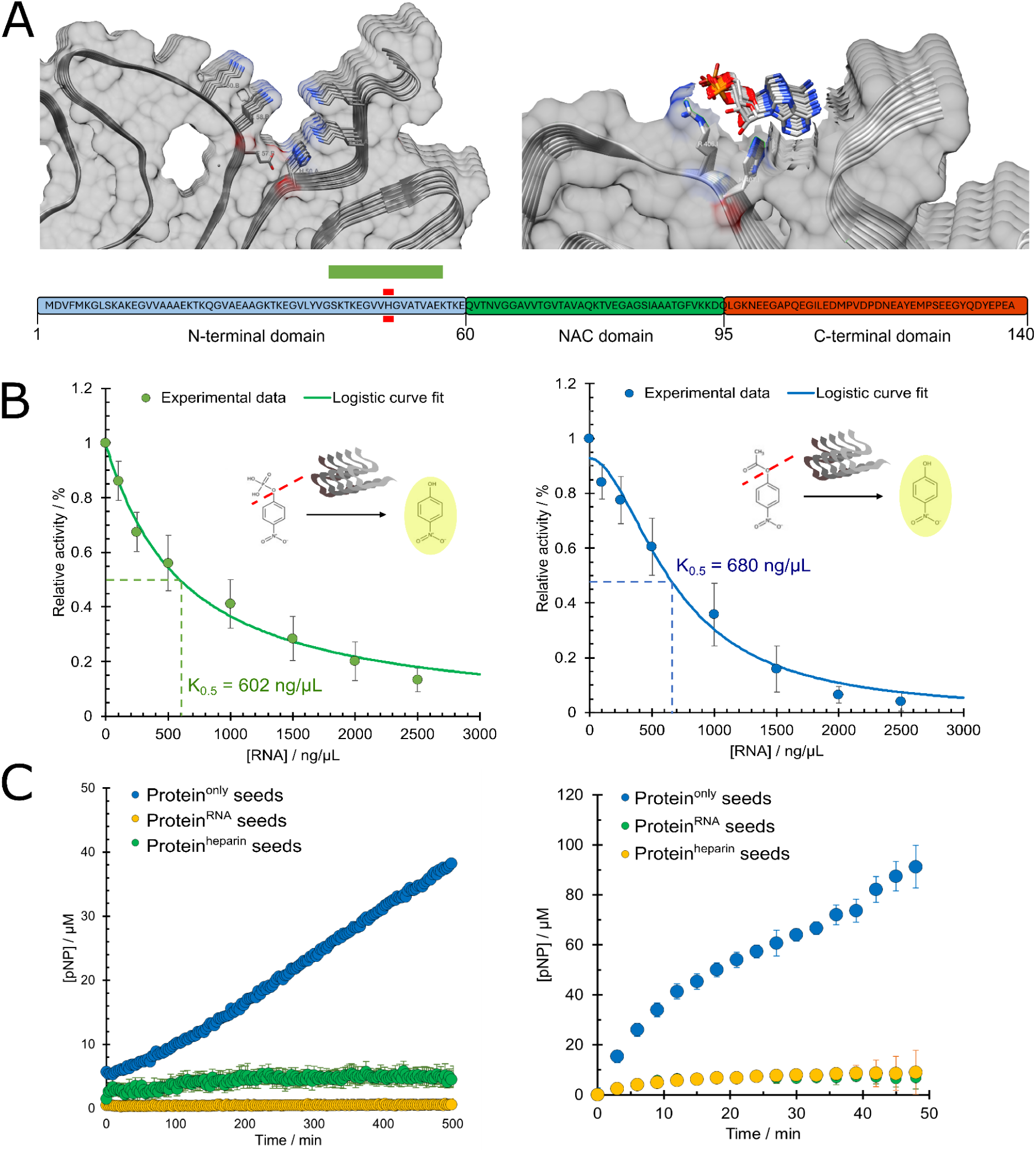
RNA can inhibit the catalytic activity of aS aggregated species. (***A***) The enzymatic activity of aS aggregated species, as reported by Horvath and colleagues, is principally linked to the H50 residue of aS (red band), indicating the residue is likely solvent-accessible in catalytically active aS aggregates. H50 also lies in the region T44 - T59 with the highest predicted RNA-binding propensity by *cat*RAPID (score > 0.5, green band, also see **Supp.** Fig. 4). An example of a structure with solvent-accessible H50 is the structure of rod 1a polymorph aS fibril (PDB 6CU7), where the arrangement of aminoacid residues around the longitudinal repeats of His50 shows a negatively (E57) and positively (K43) charged residue capable of either protonation or deprotonation of the histidine residues, thus creating a putative active site. The close spatial arrangement of histidine and arginine residues shows a degree of similarity to that of RNA-induced tau fibrils with a poly-G molecule docked during analysis (PDB 7SP1). (*B*) RNA additions incrementally inhibit the esterase and phosphatase activity of aS fibrils. Plotting the initial velocities against the concentration of RNA and fitting the data with a logistic equation (see Materials and Methods) allows the estimation of the RNA-binding affinity of the fibrils as the RNA concentration at half the maximum signal. Comparing the results obtained by measuring the esterase and phosphatase activity gives an apparent affinity constant K_0.5_ of 680 ng/μL and 602 ng/μL respectively. (*C*) Fibrils seeded by seeds grown in the presence of either RNA or heparin show almost negligent catalytic activity towards pNPA and pNPP, indicating that RNA effectively alters the structure of aS fibrils or at least the chemical environment of the catalytic amino acid residues.

We hypothesized that measuring the catalytic activity in protein^only^ and protein^RNA^ aS fibrils could both demonstrate RNA binding and confirm RNA-induced conformational changes. After validating the catalytic activity of aS fibrils (**Supp. Fig. 5, Supp. Table. 1**), we performed enzymatic assays in the presence of increasing RNA concentration. We used two substrates—para-nitrophenyl acetate (pNPA) and para-nitrophenyl phosphate (pNPP)—to avoid substrate bias. RNA addition at constant fibril and substrate concentrations markedly reduced activity towards both substrates (**Fig. 5B**). To estimate RNA’s binding affinity, we fitted the data using a four-parameter logistic equation, commonly applied when the inhibition mechanism is complex or poorly defined (20). This approach yielded apparent K_0,5_ values of 680 ng/μL (pNPA) and 602 ng/μL (pNPP). Since the assay was performed with total yeast RNA, which mostly consists of around 280 nucleotide-long fragments (BioAnalyzer analysis, data not shown), the approximate molar affinity constant could be estimated at around 7 μM. While these values are approximate due to unknown actual RNA molarity, they unequivocally indicate direct RNA-fibril interaction. When we repeated the assay with fibrils grown from protein^RNA^ seeds—but in the absence of RNA— we observed no enzymatic activity. Hydrolysis rates of both pNPA and pNPP matched baseline levels (**Fig. 5C**). These fibrils were not exposed to RNA during growth, suggesting that structural changes induced by RNA co-aggregation impair catalytic function, likely by restricting access to residues such as His50. Similarly, fibrils prepared in the presence of heparin—a known remodeler of aS fibrils—also showed no catalytic activity (**Fig. 5C**).

### aS fibrils preferentially sequester uridine-rich RNA

To investigate which RNA species interact specifically with aS fibrils, we performed RNA sequencing on RNA extracted from fibril-RNA complexes. We used total RNA from HEK293T cells instead of yeast RNA to enable efficient library preparation. In order to compare the binding preferences of aggregates, formed in the presence and absence of RNA, we performed the sequencing of both RNA extracted from fibrils, grown from monomeric aS in the presence of RNA (protein^RNA^ fibrils), as well as RNA incubated with pre-formed protein^only^ fibrils. We also performed RNA-only control experiments in parallel to allow for a differential enrichment analysis, comparing binding preferences between the two structurally distinct fibril populations (see **Materials and Methods**).

Analysis of enriched hexanucleotide motifs revealed sample-specific enrichment. Protein^RNA^ fibrils showed strong enrichment for AU-rich sequences, particularly uridine, both when controlling for transcript abundance (**Fig. 6A**) and transcript length (**Supp. Fig. 7**). In contrast, GC-rich motifs were depleted in the protein^RNA^ fibrils. To explain these findings, we turned to structural data from disease-associated aS fibrils. Notably, structures of aS amyloids from brains of patients with MSA contain unassigned electron densities closely associated with the fibrils (PDB 6XYP and 6XYO (7)). These structures feature a hydrophilic central cavity that appears to accommodate an unidentified ligand, potentially RNA, in direct contact with residues such as Lys43, Lys45 and His50.

**Figure 6.**
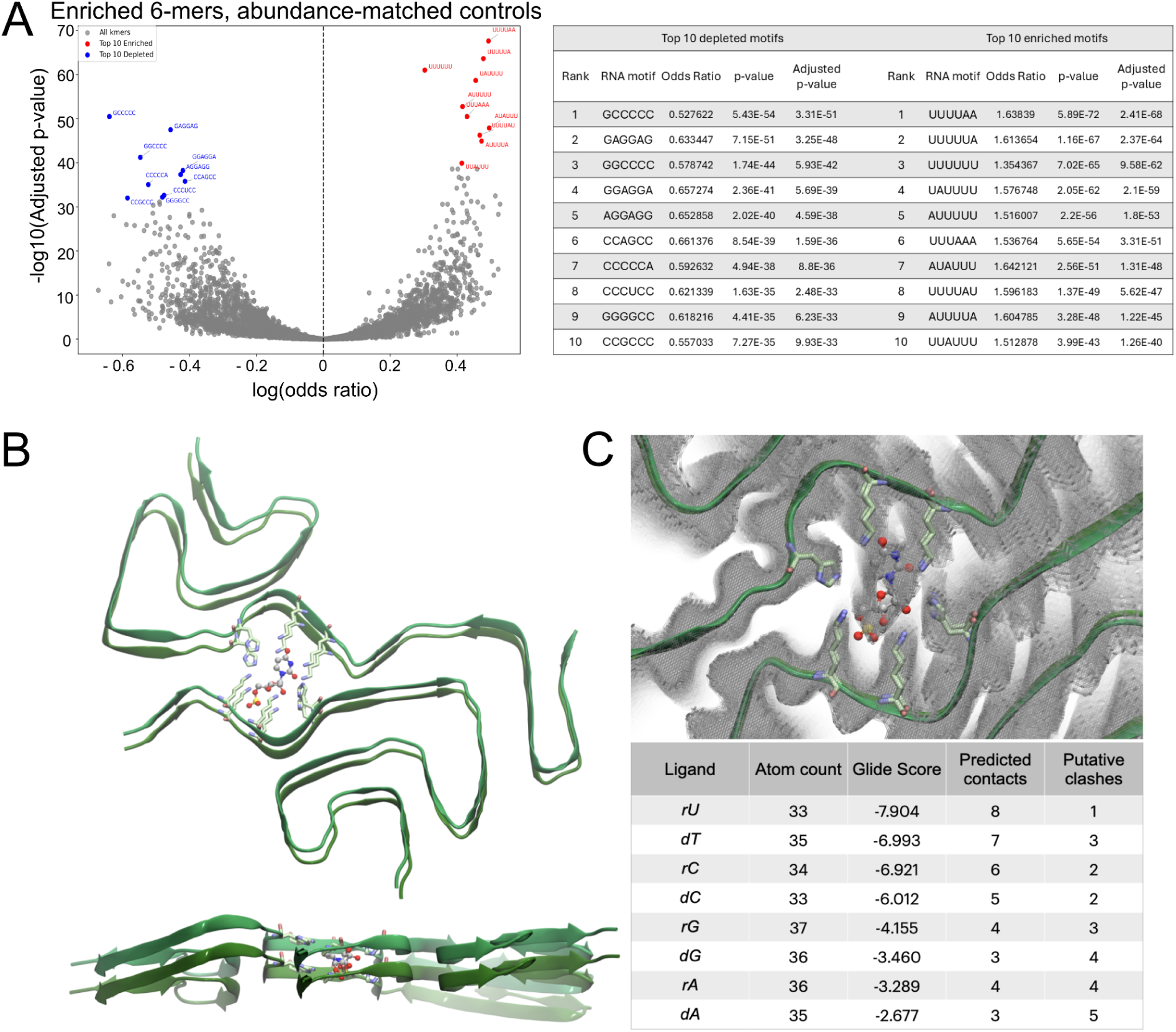
Sequencing results of the extracted RNA from aS fibrils upon coaggregation. (A) Results of the hexamer enrichment analysis comparing human transcripts specifically bound by Protein^RNA^ fibrils with an abundance-matched control set of RNAs (see Materials and Methods). Left panel: Volcano plot showing the log(Odds Ratio) versus –log₁₀*(adjusted p-value). The 10 most enriched and 10 most depleted hexamers, selected based on adjusted p-value, are highlighted in red and blue, respectively. Right panel: Table reporting the sequence, Odds Ratio, p-value, and adjusted p-value for these 20 hexamers. Motif enrichment was assessed using the chi-square test, and p-values were corrected for multiple testing using the Benjamini–Hochberg procedure. (B,C)* The structure of aS fibrils, extracted from MSA patient brain (PDB 6XYP (7) with a uridine monophosphate nucleotide docked in the central cavity show a planar orientation between the two fibril layers. Both RNA and DNA nucleotides can be docked into the unknown density (grey shaded in D) at the core of the filament, with pyrimidine nucleotides preferred likely due to spatial constraints.

To test this, taking advantage of the 6XYP cryo-EM structure (7) that paved the way for a structure-based investigation, we conducted molecular docking studies using either ribonucleotides or deoxynucleotides as putative ligands for one or two protein dimers extracted from the aS filament (**Fig. 6C-E**). The following criteria were adopted in order to filter the docking poses: (i) The positively charged sidechains were oriented towards the inner part of the site; (ii) An effective interaction between one of the two histidines shaping the binding site and the nucleotide sugar had to be present; (iii) RNA bases were oriented in planarly to the filament layering in order to let a good positioning of the phosphate group and allow the presence of phosphodiester bonds with the previous and the following nucleotide forming the negatively charged phosphate backbone of a putative RNA or DNA strand occupying the unassigned electron density. Pyrimidine nucleotides (uridine and cytidine), due to their smaller size, fit well into the planar pose of the fibril structure (**Fig. 6B**). In contrast, purines (adenine and guanine) could only bind in interplanar pockets between adjacent monomers (**Fig. 6C**), leading to lower score predictions. Binding was further influenced by the electrostatic environment: the cavity contains positively charged residues (two histidines and four lysines for each dimer/layer), favouring nucleotides with electron-donor atoms capable of hydrogen bonding. Moreover, as an indirect consequence of (ii) criterion, ribonucleotides were preferred over deoxynucleotides, likely due to their smaller size and additional hydroxyl group, which may enable better interactions (with histidines) despite some steric costs. These docking results are consistent with our earlier finding that aS also sequesters DNA, though less efficiently than RNA (5).

Together, these data suggest that nucleic acids could fit within the unidentified electron-dense cavities observed in patient-derived aS structures. This provides a plausible structural basis for the sample-specific enrichment observed in the sequencing results and highlights a molecular mechanism by which aS fibrils selectively sequester RNA based on both sequence and structure.

## Discussion

Our study uncovers a multifaceted role for RNA in modulating the aggregation, propagation, and structural diversification of aS fibrils. Building upon prior findings implicating nucleic acids as cofactors in amyloidogenesis (33, 34), we show that RNA not only influences the kinetics of aS aggregation but also decisively shapes the structural and functional properties of resulting amyloid assemblies. This work refines the emerging paradigm that amyloid fibrils are not merely proteinaceous but may co-assemble with endogenous molecules—particularly RNA—giving rise to distinct, functionally diverse polymorphs.

Extending our previous findings that RNA accelerates the *de novo* aggregation of aS (5), in this work we demonstrate that RNA co-aggregation alters aS seeding efficacy in a concentration- and seed-dependent manner. Under seed-saturating, quiescent conditions that minimize mechanical fragmentation, protein^only^ seeds exhibit faster growth than their RNA-containing counterparts. This suggests that RNA co-incorporation during aggregation induces an altered fibril conformation which sterically impairs monomer addition.

The addition of RNA to protein^only^ seeds reveals a dual effect. At high seed concentrations (5–10 μM), it enhances elongation when added to protein^only^ seeds, yet it suppresses fibril growth at lower concentrations. Furthermore, it alters the kinetic path by promoting elongation and suppressing secondary nucleation. This is highly relevant considering the predominance of secondary nucleation in physiological settings and its central role in toxic oligomer formation (35). Secondary nucleation has been shown to occur via transient interactions between the fuzzy coat of aS fibrils and the monomer with an estimated K_D_ of around 1 mM (36), implying that fibril-RNA binding with a higher affinity could potentially impair this mechanism.

Even more strikingly, RNA induces the efficient elongation of protein^RNA^ seeds with kinetics on par with protein^only^ seeds. This directly supports the hypothesis that RNA-induced fibrils adopt distinct conformations that require continued cofactor interaction—a phenomenon mirrored in tau and MSA-associated fibrils (4, 7). This behavior hints at a conformational gating mechanism or competitive binding equilibrium, contingent on stoichiometry and fibril morphology, where interactions between monomeric aS and fibril-bound RNA may either facilitate or restrict H-bond formation between the fibril end and monomeric protein to facilitate templated fibril elongation (37, 38).

Spectroscopic and morphological analyses further reveal that RNA co-aggregation promotes the formation of more compact and rigid fibrils due to a stronger network of hydrogen bonds that stabilize the structure, as evidenced by cross-β shifts in FTIR and distinct height profiles in previous AFM analysis (34) . These RNA-containing fibrils differ significantly from their protein^only^ counterparts, exhibiting lower heterogeneity, a feature previously linked to altered seeding potency and cellular toxicity (9, 34). However, whether RNA-induced polymorphs confer differential pathogenicity *in vivo* remains to be determined, especially given the structurally different patient-derived amyloid polymorphs (7, 39). A strong implication could be the RNA-conferred resistance to proteolytic degradation. While

RNase-treatment can rescue the impeded pK degradation of protein^only^ fibrils with RNA bound to the fibril surface, structural RNA could render heterotypic protein^RNA^ fibrils much more resistant to cellular clearance mechanisms. Furthermore, RNA interaction with positively charged amino acids like lysine on aggregate surface could potentially impede ubiquitination and chaperone recognition (40), allowing aggregates to evade cellular degradation and thereby fostering their accumulation and persistence.

One of the most compelling findings of our study is the characterisation of the direct interaction between RNA and aS fibrils by means of inhibition of fibril enzymatic activity, centered around His50. The estimated affinity of binding would be more that two orders of magnitude lower than the estimated affinity between aS monomers and fibrils (36), making it potentially physiologically relevant. Similarly, analysis of the proposed inhibition mechanism reveals a complex scenario with multiple independent binding sites on the fibril (**Supp. Fig. 6**). Conceptually, this fits well a scenario with multiple RNA molecules binding the fibril at multiple sites with different affinities based on their molecular properties. Furthermore, the loss of catalytic activity in RNA- or heparin-induced fibrils indicates a change in the chemical environment of His50, rendering it inaccessible to the solvent. This result is noteworthy as His50 is buried in the fibril core in the two structures of aS amyloids isolated from brains of patients with Lewy-body dementia and MSA (7, 39). These disease-associated structures have yet to be reproduced *in vitro* and our findings suggest RNA may be a key factor in directing the assembly of disease-relevant aS fibril conformations.

Structural modeling supports this finding, showing that pyrimidine nucleotides engage hydrophilic pockets and positively charged residues (including His50) within patient-derived MSA fibrils. These interactions appear to be driven by electrostatic complementarity rather than random sequestration, suggesting that specific RNA motifs may selectively template or stabilize particular fibril conformations. While these insights remain preliminary and warrant validation through targeted mutagenesis and biochemical binding assays. RNA sequencing analysis however supports this claim partially, as RNA extracted from coaggregated protein^RNA^ fibrils appear to be enriched in U-containing motifs. While our earlier work predicted that aS preferentially interacts with GC-rich elements (41), and the recent study by Matsuo et al. (6) confirmed that RNA G-quadruplexes can directly bind and promote aS aggregation *in vitro* and in neurons, our current findings point to a different subset of RNAs becoming embedded within mature aggregates. Specifically, we find that AU-rich, uridine-containing transcripts are significantly enriched in the coaggregated RNA fraction. This suggests a mechanistic distinction between RNAs that initiate aggregation, such as structured, G-rich RNAs capable of promoting phase transitions, and those that are retained within the fibrillar structure. Such decoupling supports a two-stage model: GC-rich RNAs may act transiently as nucleating cofactors, while AU-rich sequences are more stably sequestered during fibril elongation. This separation of function is in line with the broader view that RNA modulates both the kinetics and structural evolution of amyloids in distinct, temporally defined ways.

In conclusion, our findings establish RNA as a versatile and dynamic cofactor in aS aggregation. Far from being a passive bystander, RNA actively shapes the structural, kinetic, and functional attributes of aS fibrils. The possibility that RNA could differentially propagate or exclude certain amyloid strains—such as those associated with Lewy body dementia versus MSA—adds a critical layer to our understanding of strain heterogeneity. This work not only redefines our understanding of amyloid biology but also opens new avenues for therapeutic intervention—potentially by targeting RNA-fibril interactions to destabilize pathogenic polymorphs or enhance aggregate clearance. Further investigation into RNA identity, binding motifs, and phase-separated environments will be essential to fully elucidate RNA’s role in the pathogenesis of synucleinopathies.

## Materials and Methods

### Preparation of alpha-synuclein seeds

The expression and purification of recombinant aS was performed as described previously (5). Seeds were prepared by aggregating 100 μM aS in the presence and absence of 500 ng/μL total yeast RNA (totRNA, Roche, cat. no. 10109223001) on black, clear-bottom 96-well plates (VWR, cat. no. VWRI732-3738) in the presence of a borosilicate glass bead with a 3 mm diameter. The samples were let to aggregate at 37 °C with constant shaking in a Spark microplate reader (Tecan) for 24 h. The samples were then transferred into Protein LoBind tubes (Eppendorf, cat. no. 0030108116) and centrifuged at 18.000 g for 1 h at room temperature. The supernatants were removed and the concentration of soluble protein was determined via bicinchoninic acid assay (BCA) assay (Thermo Fisher Scientific, cat. no. 23225) or Qubit Protein assay (Thermo Fisher Scientific, cat. no. Q33212). The seed concentration was determined by subtracting the concentration of the soluble protein in the supernatant from the initial protein concentration. The pelleted aggregates were resuspended in sterile, 1xPBS and centrifuged again at 18.000 g for 1 h at room temperature. The supernatant was discarded and the pellet was resuspended in sterile, 1xPBS and RNase A (Thermo Fisher Scientific, cat. no. EN0531) was added to a final concentration of 50 μg/ml in order to degrade any sequestered RNA. The reaction mixtures were incubated at 37 °C for 1 h and then 1 U/ml RiboLock RNase inhibitor (Thermo Fisher Scientific, cat. no. EO0381) was added to inhibit the action of RNase A (Thermo Fisher Scientific, cat. no. EO0381). The samples were centrifuged at 18.000 g for 30 min at room temperature and the supernatant was discarded. This washing step was repeated twice, afterwards the pellet was resuspended in sterile, 1xPBS, aliquoted in Protein LoBind tubes, flash-frozen, and stored at - 80 °C.

### Seeded aggregation assays

A fresh aS aliquote was quickly thawed and filtered through a 0.22 μm pore syringe filter (Merck - Sigma Aldrich, cat. no. SLGVR04). Total yeast RNA powder (Roche, cat. no. 10109223001) was resuspended in aggregation buffer (20 mM potassium phosphate pH 7.2, 100 mM KCl, 5 mM MgCl_2_) and its concentration was measured via NanoDrop (Thermo Fisher Scientific, cat. no. ND-ONE-W). An aliquot of seeds was quickly defrosted and let to sonicate in a water bath sonicator for 10 min at room temperature. All the reagents were mixed in Protein LoBind tubes and Proteostat^TM^ (Enzo Life Science, cat. no. ENZ-51023-KP002) was added as a fluorescent aggregation marker. The final concentration of monomeric aS stock was 50 μM and 500 ng/μL for totRNA in aggregation buffer, while the final concentration of the seeds ranged between 1 μm and 10 μm. Aliquots of 100 μL were transferred to wells in black, clear-bottom 96-well plates (VWR, cat. no. VWRI732-3738), sealed and let to aggregate at 37 °C without agitation until fluorescence reached a plateau (ca. 40 h). Proteostat^TM^ fluorescence was recorded at the bottom of wells every 15 min and every condition (e.g. seed concentration) was represented in at least four replicates.

### Analysis of aggregation assay data

As fluorescence data was recorded in 5 different points in the well, the signal was first averaged at all timepoints. Background (buffer or buffer/RNA) was subtracted from the samples and fluorescent traces were then normalized according to minimal and maximum well fluorescence read. The reads at t < 1 h were then fitted with a linear trendline and the gradient of the slope was recorded for each individual replicate. These were then averaged for every condition and represented as the aggregation rate.

In order to determine the fluorescence trace curvature, the data points at t > 10 h were fitted to a sigmoid equation as described previously (5). The coefficient h was calculated from the individual fits and averaged across all conditions and represents the curvature of the curve, indicating the presence of secondary processes.

### Proteinase K fibril degradation assay

The fibrils were prepared as for the enzyme catalysis reactions (see below). After centrifugation, the pellet was discarded and the pelleted aggregates were resuspended in 1x PBS with 0.5 μg/mL pK (Roche, cat. no. 3115887001) added. The reaction mixtures were incubated at 37 °C with mild agitation for 5 h. Afterwards, 6x SDS-PAGE loading dye was added, the samples were boiled at 95 °C for 10 min and loaded on a 4-12 % denaturing polyacrylamide gel (Invitrogen, cat. no. NP0321). After electrophoresis, the gel was stained with InstaBlue (Expedeon, cat. no. ISB1L).

### Fluorescence fibril degradation assays

After the fluorescence stabilized at the maximum value for all the conditions, the microplate with the aggregated aS was briefly removed from the plate. For every condition, samples in half of the wells were left as controls (2 μL of PBS was added directly to each well) and in the other half, RNase A treatment was performed by adding 2 μL of stock RNase A (Thermo Fisher Scientific, cat. no. EN0531) in PBS to every well. The microplate was immediately resealed and returned to the microplate reader preset to 37°C. Fluorescence was read every 5 min at the bottom of the wells, with 10 s of shaking before each measurement, for 5 h. The microplate was then again removed from the plate reader and 2 μL of 2.5 mg/mL pK (Roche, cat. no. 3115887001) solution in PBS was added to each well (final concentration 0.5 μg/mL). The plate was again immediately resealed and returned to the microplate reader preset to 37 °C. Fluorescence was measured every 5 min at the bottom of the wells, with 10 s of shaking before each measurement, for 20 h.

### Analysis of fibril degradation assays

The fluorescence signal was recorded in 5 different points in the well and was first averaged at all timepoints. Background (buffer or buffer/RNA) was subtracted from the samples and fluorescent traces were then normalized according to minimal and maximum well fluorescence signal. Fluorescence traces were linearised and fitted to the model function (1):

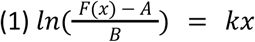

with *A* and *B* as the fitting parameters. The parameter *k* or the slope of the initial is the linear part of the fluorescence decay, representing the initial reaction rate. The initial rates for every condition were averaged and compared using the two-way paired Student t-test.

### Fibril enzymatic activity assay

The protocol for fibril enzymatic activity was used as described by Horvath and colleagues (16). The fibrils were prepared by mixing 200 μM monomeric aS with 5 μM seeds, prepared as described above, in aggregation buffer in Protein LoBind tubes. Heparin seeds were prepared according to the above-described protocol, with totRNA substituted for heparin (Merck - Sigma Aldrich, cat. no. H3149). The fibrils were grown at 37 °C degrees without agitation for 5 days. The fibrils were always prepared fresh on the day of the enzymatic assay. The samples incubated at 37 °C were centrifuged at 18.000 g for 1 h at room temperature and the protein concentration of the supernatant was determined with a BCA assay. The fibril concentration was then determined by subtracting the supernatant concentration from the initial monomer concentration (200 μM). The pellet after centrifugation was resuspended with sterile 1xPBS to a monomer-equivalent concentration of 200 μM. Reaction mixes were prepared by addition of sterile 1xPBS, 0.5 M MgCl_2_ and freshly resuspended totRNA to Protein LoBind tubes. Fibrils were sonicated in the water bath for 15 min at room temperature and the sonicator setting of 8, then added into the reaction mixes. To stimulate fibril-RNA interaction, the reaction mixes were incubated at room temperature with gentle mixing for 30 min. The substrate was then added to a final concentration of 10 mM (pNPP, Merck - Sigma Aldrich cat. no. 71768) or 5 mM (pNPA, Merck - Sigma Aldrich cat. no. N8130). The reaction mixture was briefly mixed with a pipette and aliquoted into Costar transparent 96-well plates (Corning, cat. no. 3370). The plates were sealed and incubated at 37 °C for up to 16 h. The reactions were followed by measuring the absorbance at 410 nm (A_410_) every 3 min in a Spark multiplate reader (Tecan).

### Analysis of enzymatic activity assays

The time-resolved A_410_ data from the enzymatic assays were first blanked by subtracting the A_410_ values from wells containing only substrate and buffer. This way, any catalytic activity due to potential enzymatic contaminants in the buffer, as well as auto-hydrolysis of the substrate, was eliminated. The A_410_ values were then converted into the molar concentration of the product 4-nitrophenol (pNP) according to the compound’s extinction coefficient of 7500 M^−1^cm^−1^ measured in 50 mM phosphate buffer at pH 7.

The basic kinetic parameters such as K_M_, k_cat_ and ε were determined as outlined by Horvath and colleagues (16). For the determination of the binding affinity of RNA, the fitted v_0_ at different concentrations of RNA were plotted against the initial RNA concentration. The resulting values were fitted by least-square linear regression with a logistic curve with four parameters (2):

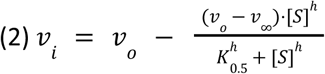

with v_0_, v_∞_, K_0.5_ and h as the fitting parameters and [S] the initial substrate concentration.

The estimation of the inhibition mechanism was done via the Lineweaver-Burk and Dixon plots. Lineweaver-Burk plot is a linearised version of the Michaelis-Menten plot, with 1/v plotted against 1/[S]. The intersections of the data with the x and y axes can identify the competitive character of the inhibition mechanism, however they cannot distinguish between linear and nonlinear mechanisms. This was done using the shape of the curve in the Dixon plot, with 1/v plotted against the concentration of the inhibitor. A linear shape indicates a linear inhibition mechanism, while an exponential curve can denote other mechanisms such as a hyperbolic or parabolic inhibition.

### Infrared spectroscopy of aS fibrils

#### FTIR measurements

For the IR experiments, 20 μL were deposited at the center of CaF_2_ windows and let to dry completely under a gentle nitrogen flow in low-humidity conditions. FTIR spectra were recorded with a Bruker VERTEX 70v spectrometer (Bruker Optic GmbH, Ettlingen, Germany) equipped with a N_2_-cooled mercury-cadmium-telluride (MCT) detector from Infrared Associates Inc. The interferometer was operated in low vacuum conditions to remove the IR absorption of atmospheric CO_2_ and water vapor. Spectra were recorded in the range from 1200 to 6000 cm^-1^ with a resolution of 2 cm^-1^ and 512 scans.

#### Micro-FTIR measurements

Micro-FTIR spectra were acquired using a Vertex 80 spectrometer (Bruker Optic GmbH, Ettlingen, Germany) combined with a Hyperion 3000 IR-microscope (Bruker Optic GmbH, Ettlingen, Germany) at the IRIS beamline at BESSY II, Helmholtz Zentrum Berlin. The Hyperion 3000 microscope, continuously purged with gaseous nitrogen, was equipped with a 15x Cassegrain objective lens (numerical aperture, NA = 0.4). Data was acquired using a liquid-nitrogen cooled 64 x 64 pixel focal plane array (FPA) detector, covering a total area of 173 x 173 μm^2^ (single pixel area is 2.7 x 2.7 μm^2^). Spectra were recorded in transmission mode using the internal Globar source, within the spectral range 800-4000 cm^-1^ with a resolution of 2 cm^-1^ and 64 scans.

#### Analysis of IR spectra

FTIR spectra were baseline-corrected, spline-smoothed, and normalized in IgorPro 6.22A, which was also used to compute second-order derivatives. For micro-FTIR ω_cross–β_ analysis of protein^only^ and protein^RNA^ fibrils (FPA detector), spectra (64 × 64 per acquisition) were baseline-corrected over 1480–1730 cm⁻¹ and integrated over 1490–1720 cm⁻¹ to generate amide-I/II intensity maps in OPUS 8.7. Second-order derivatives were calculated using the Savitzky–Golay method (9-point smoothing), and an ω_cross–β_ map was generated by identifying derivative minima within 1624–1632 cm⁻¹ using the *peak picking* command in OPUS software. Spectra lacking a single minimum in this range were excluded for low signal-to-noise ratio. To remove pixels with low protein content, an intensity mask (50–100% of maximal amide-I/II signal) was generated in the amide-I/II map in Gwyddion 2.61 and applied to the ω_cross–β_ map. The masked pixels were then used to generate the histograms of the ωcross–β distribution in IgorPro 6.22A..

#### AFM topographic map

Three aliquots of 2 μL of protein^only^ seeds and protein^RNA^ seed samples were drop-cast onto 1 cm x 1 cm template-stripped gold surfaces (Platypus Technologies, 0.3 nm nominal roughness) for 5 min, then rinsed with 40 μL Milli-Q water to achieve a sparse distribution of deposited aS fibrils. The samples were let to dry for 1 h under a gentle nitrogen flow in a controlled low-humidity environment. AFM topographies of the fibril bundles were measured using the AFM module of the neaSNOM system (attocube systems AG, Germany) at the IRIS beamline at BESSY II, Helmholtz Zentrum Berlin. For the protein^only^ seeds, an Arrow-NCPt (tip radius <25 nm) Pt/Ir-coated AFM probe (Nanoworld AG, Switzerland) was used, while for the protein^RNA^ seeds, a PPP-NCSTAu (tip radius <50 nm) Au-coated AFM probe (Nanosensors, Switzerland) was employed. Measurements were performed in tapping mode at room temperature.

#### RNA extraction from aS aggregates

RNA was extracted from HEK293T cells, grown in culture according to standard procedures, with the Directo-Zol RNA Mini-Prep kit (Zymo #R2050). DNase-treatment was performed for all the samples before the final step of the purification. Total cell RNA was eluted using nuclease-free water. 50 μM monomeric aS was aggregated in aggregation buffer both in the presence and absence of 500 ng/μL total cell RNA in 1,5 mL DNA LoBind tubes (Eppendorf) at 37 °C with constant agitation for 72 h. 500 ng/μL total cell RNA in aggregation buffer was coincubated as a control. The samples were centrifuged for 1 h,

18.000 g at 4 °C. For protein^RNA^ fibrils, the supernatants were removed and preserved as the “non-sequestered fraction”. For the protein^only^ fibrils, supernatants were removed and pellets were resuspended in the same volume of aggregation buffer with 500 ng/μL total cell RNA. These samples, termed “RNA pull-down”, were subjected to incubation with slight agitation at room temperature for 2 h. Again, 500 ng/μL total cell RNA in aggregation buffer was coincubated as a control. Afterwards, the samples were centrifuged for 1 h, 18.000 g at 4 °C, and the supernatants were removed and preserved as the “non-sequestered” fraction. The pellets were washed 2 times with the same volume of aggregation buffer at 4 °C. Afterwards, the pellets were gently resuspended in aggregation buffer and centrifuged again for 1 h, 18.000 g at 4 °C. RNA was extracted from the pellets as described before (5).

The experiment was repeated 3 times independently and each time, 4 individual replicates for every condition (coaggregation and “pull-down”) were performed. The quality of the RNA was assessed individually with BioAnalyzer and 3 replicates for each experiment were chosen for library preparation.

#### RNA sequencing experiment and analysis

Next-generation sequencing libraries were prepared using the Illumina Total RNA Prep with Ribo-Zero Plus kit. Sequencing was conducted on a Illumina NovaSeq 6000 instrument. The Illumina bcl2fastq2 (version 2.20) utility was employed for BCL-to-FASTQ conversion, demultiplexing, adapter trimming, and filtering out trimmed reads shorter than 35 nucleotides. Reads mapping to human rRNA sequences were identified and removed using Bowtie2 (version 2.3.4.1) (42) . The remaining reads were aligned to the human genome (hg38) with STAR (version 2.7.6a) (43), leveraging the GENCODE release 43 GTF file for splice junction annotation. The alignments were performed with parameters *--quantMode GeneCounts --outSAMstrandField intronMotif --outSAMtype BAM SortedByCoordinate --readFilesCommand zcat --peOverlapNbasesMin 10*. The resulting alignment files were merged into a single file using SAMtools (version 1.7) (44) with the merge command. The merged BAM file served as input for transcriptome assembly with StringTie (version 2.2.1) (45), using the *--rf* option and the GENCODE GTF file as a reference annotation. Post-assembly, the GTF file generated by StringTie was processed with Python 3.7 to retain only transcripts assigned to known genes while excluding rRNA annotations. Further filtering was performed to retain the most relevant transcripts per gene based on expression levels (FPKM) and genomic uniqueness. Transcripts were prioritized by FPKM, and redundancy was assessed by comparing nucleotide coordinate overlaps with higher-expressed isoforms. Transcripts were excluded if their unique nucleotide regions (i.e., regions not shared with higher-priority transcripts) were <50 bp or their FPKM was <10% of the top isoform. Gene coordinates were updated to reflect the retained transcripts, and a reduced GTF file was generated for downstream analyses. Genomic coordinates of multi-exon genes were written to a BED file, and the BEDTools getfasta utility (46) was used to extract precursor transcript sequences. The GffRead utility (47) was employed to retrieve transcripts sequences from the reduced GTF file. The two sequence sets were combined to generate a reference transcriptome. Salmon (version 1.3.0) (48) was then employed to quantify transcript abundances for each sample against this reference transcriptome, using parameters *-l ISR --gcBias --numBootstraps 30 –validateMappings*. The prepare_fish_for_sleuth function from the wasabi (version 1.0.1) R package (available at https://github.com/COMBINE-lab/wasabi) was employed to convert Salmon output into sleuth-compatible format. The tximport function from the tximport (version 1.18.0) (49) R package was used to import transcript-level abundance estimates into an R (version 3.6.3) object. Differential transcript abundance analyses were performed using the sleuth (version 0.30.0) (50) R package, considering only transcripts with average FPKM in input samples greater than 0.1. The sleuth_wt function was used to perform a Wald test identifying transcripts enriched in the protein^RNA^ or protein^only^ samples with respect to their RNA pull-down controls – i.e. those transcripts which are bound by protein^RNA^ or protein^only^ fibrils (protein^RNA^-bound and protein^only^-bound, respectively), selected based on a q-value threshold of 0.05 and a log_2_(fold-change) threshold of 0.58. To identify transcripts with different fibril/control enrichment between protein^RNA^ and protein^only^ experiments, we performed a likelihood ratio test (LRT) using the sleuth_lrt function with a full model and a reduced model. Transcripts preferentially bound by the protein^RNA^ fibrils with respect to the protein^only^ fibrils were identified as the subset of protein^RNA^-bound ones having an LRT q-value < 0.05 and a fibril/control ratio in the protein^RNA^ experiment at least 1.5 times greater than the one observed in the protein^only^ experiment (protein^RNA^-enriched transcripts).

To assess the sequence properties of protein^RNA^-enriched transcript in terms of 6-mer enrichment, we compared them against two different control sets of transcripts. These sets were generated by selecting, for each protein^RNA^-enriched transcript, the RNA with the same type and with the most similar abundance (abundance-matched controls) or length (length-matched controls) among those having an LRT q-value > 0.25. This approach ensured that the comparison accounted for differences in transcript abundance or length, enabling a more robust analysis of 6-mer enrichment specific to the protein^RNA^-enriched transcripts. Only non-precursor molecules were included in the analysis. For each 6-mer present in the whole sequence set, a chi-square test was conducted to evaluate differences in proportional representation between protein^RNA^-enriched transcripts and the abundance-matched or length-matched control set. P-values were adjusted for multiple testing using the Benjamini-Hochberg correction.

#### RNA and DNA computational docking

The deoxyribo- and ribonucleotides monophosphate used as ligand molecules in this *in silico* procedure were retrieved from the PDB DB (http://RCSB.org) and imported in Maestro [*Schrödinger Release 2023-2: Maestro*; Schrödinger LLC: New York, NY, USA, 2023]: first, the ligand preparation was performed by means of the LigPrep module, with the prevalent protonation and tautomeric state assignment; whereas the Protein Preparation Wizard was used to make the set up of the protein models for the aS fibril; the latter was *ad hoc* trimmed (starting from the PDB id 6XYP structure (7), to contain one or two protein dimers only; the hydrogen assignment was based on a h-bond network optimization criteria. The docking grid center was set as the centroid of Lys43, Lys45 and His50 residues of each aS chain included in the single or the double dimer, and the box size was chosen in accordance with a maximum ligand length of 20 Å. The Glide protocol included three-steps: a SP, an XP, and an IF docking run. In a nutshell, in the SP docking the best docking poses are selected on the basis of a series of hierarchical evaluations: first a set of ligand conformations is generated by means of an extensive conformational search, then comes the assessment of the spatial compatibility between each ligand conformation and the protein. A ligand minimization in the target using the OPLS3 force field (51) is performed on the poses selected from these filters. In order to optimize the torsional angle energy, the lowest energy resulting poses are then Monte Carlo simulated, finally ranking them using the default GlideScore scoring function (52). At variance with the former, the XP protocol follows an anchor-and-grow sampling based approach and a stricter scoring function – with respect to the SP docking one (53) – evaluates each intermediate conformation. On the contrary, the IF procedure performs the docking of the ligand with both a van der Waals radii reduction and a non-bonded energy interactions cut-off increase. In order to allow an improved ligand accommodation (for each of the obtained poses), the Prime module performed a prediction step on the surrounding amino acids by re-orienting the residue side chains. Both ligand and these residues were then minimized and each ligand was re-docked (54, 55). The results, including five docking poses, were ordered according to the GlideScore, and the top pose for each deoxyribo- and ribonucleotides monophosphate was further considered for the comparative analysis.

## Acknowledgments

The authors would like to thank the RNA flagship at IIT, Istvan Horvath and Pernilla Wittung Stafshede for helpful discussions and insights. The first author would like to thank the corresponding author’s incredible patience and support in letting him procrastinate and reflect on potential implications of these results for way too long of a time. AI, MET, and VG would like to thank Helmholtz-Zentrum Berlin für Materialien und Energie for the micro-FTIR and AFM measurements carried out at the IRIS beamline of BESSY II.

## Contributions

GGT and JR conceived the study. JR performed all biochemical experiments. EZ, GGT, and JR wrote the manuscript. GGT and EZ supervised JR. AC analyzed the transcriptomic data, and FDP ran the dockings and analyzed them to generate the structural models. VG, MET, and AI performed the infrared spectroscopy experiments and analyzed infrared data with JR. All authors discussed the results, provided critical feedback, and approved the final manuscript.

