## Supplementary Figures and Materials for "RNA interactions drive structural and functional diversification of α-Synuclein fibrils"

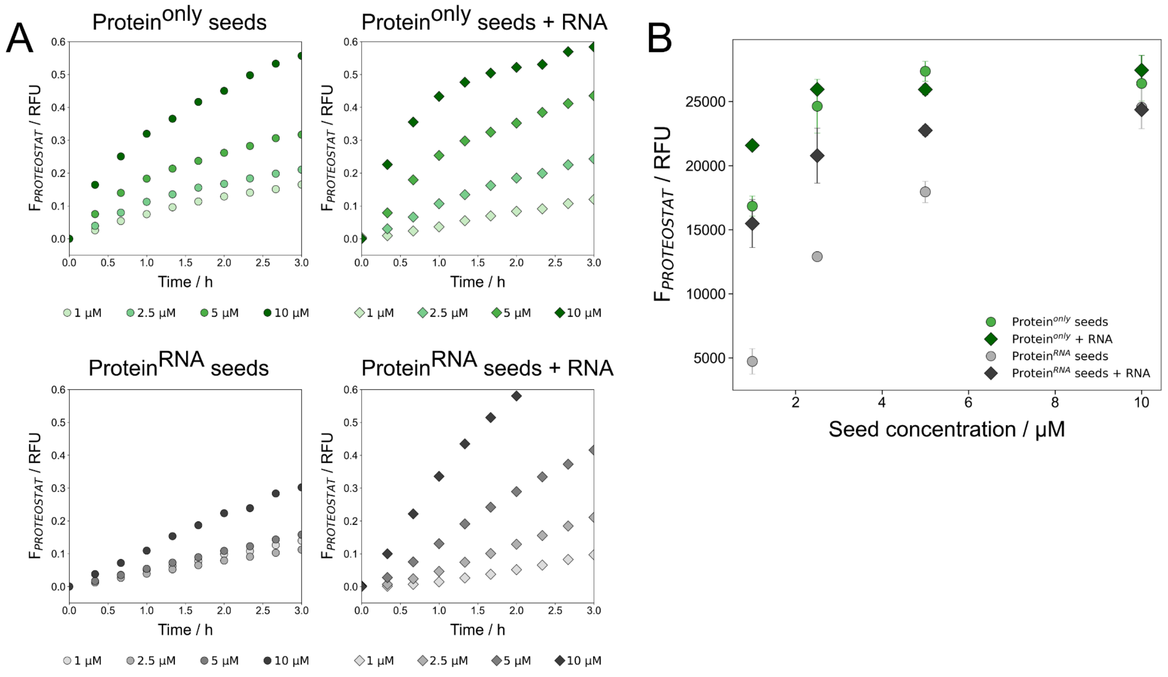


**Supplementary figure 1.** (*A*) Early-stage aggregation kinetics of 50 µM aS at varying seed concentrations and in the presence or absence of RNA. These clearly show the effect of different seed type, as well as the addition of RNA on the aggregation of aS. Protein$^{RNA}$ seeds have a much lower seeding capacity compared to protein$^{only}$ seeds and only start effectively accelerating the aggregation kinetics at 10 µM concentration. The presence of RNA significantly accelerates the kinetics in both cases. (*B*) Peak Proteostat™ fluorescence signal at varying seed concentrations. The peak fluorescence signal roughly corresponds to the amount of aggregated species, providing an estimate of the seeding ability of different seed species. Protein$^{RNA}$ seeds have the overall lowest peak fluorescence, further indicating their lower seeding ability compared to protein$^{only}$ seeds as already demonstrated by the slower aggregation kinetics.

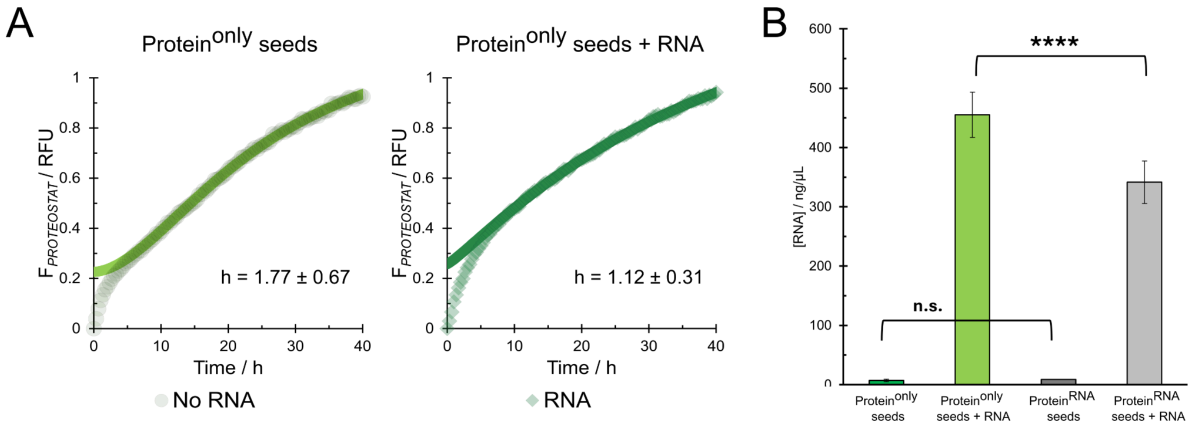


**Supplementary figure 2.** (*A*) Fitting the later exponential stage of fibril growth for protein$^{only}$-seeded kinetics with the sigmoid equation (see **Materials and Methods**) results in coefficients h of 1.12 ± 0.31 in the presence and 1.77 ± 0.67 in the absence of RNA ($p = 0.012$, $N = 8$, two-way Student t-test). This indicates an absence of secondary processes in the presence of RNA and potentially implies that while RNA increases the elongation rate, it also reduces the surface-mediated secondary nucleation processes. (*B*) Soluble RNA quantification after aggregation shows a significant decrease for reactions, seeded with protein$^{RNA}$ seeds ($p < 0.001$, $N = 8$, Student t-test).

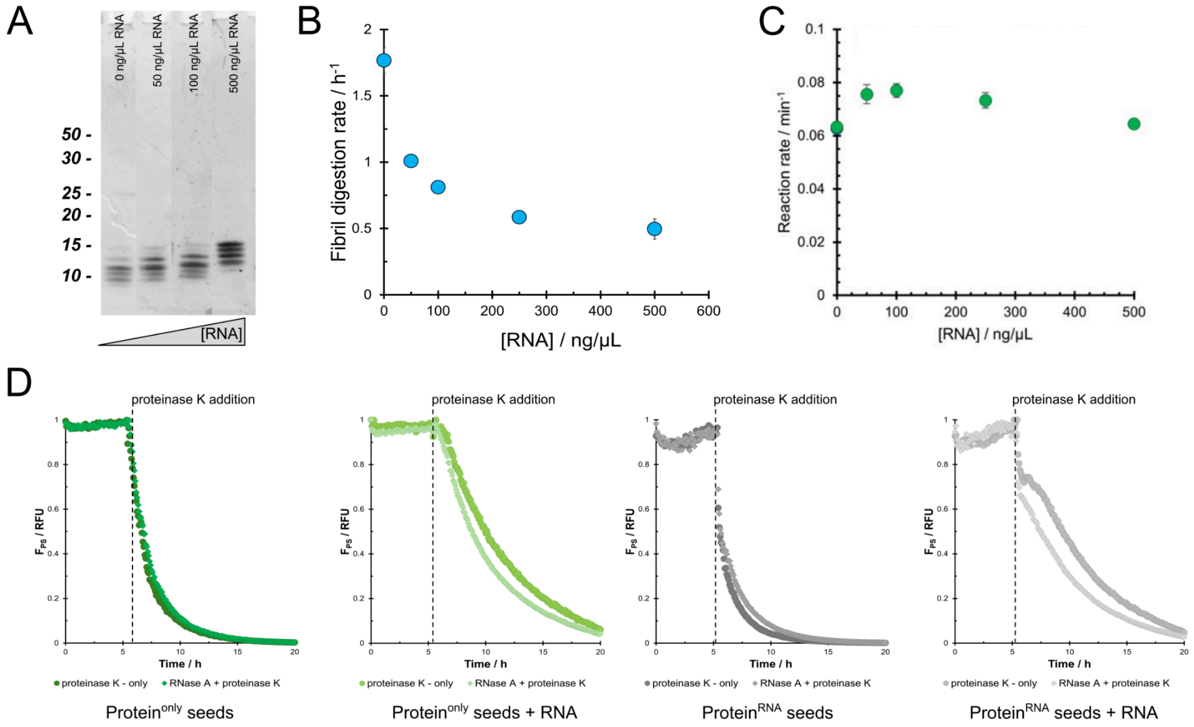


**Supplementary figure 3.** (*A*) SDS-PAGE gel analysis reveals differences in the degradation of aS aggregates by 0.5 µg/mL pK when incubated with different concentrations of RNA. (*B*) aS fibril degradation rate with pK decreases exponentially with increasing RNA concentrations. (*C*) Degradation of the enzymatic substrate pNPA by proteinase K in the presence of RNA. The rate of pK activity does not decrease at higher RNA concentrations, indicating that RNA does not inhibit pK activity directly. (*D*) Treatment of aS fibrils with RNase A for 5 h at 37 °C (lighter coloured traces) does not result in changes in fluorescence intensity with respect to controls (darker coloured traces), implying that RNase A itself does not affect the fibrils integrity. In contrast, RNase A-treated samples show a faster decay of fluorescence upon pK addition in the case of fibrils aggregated with RNA. No change is observed for fibrils, aggregated in the absence of RNA.

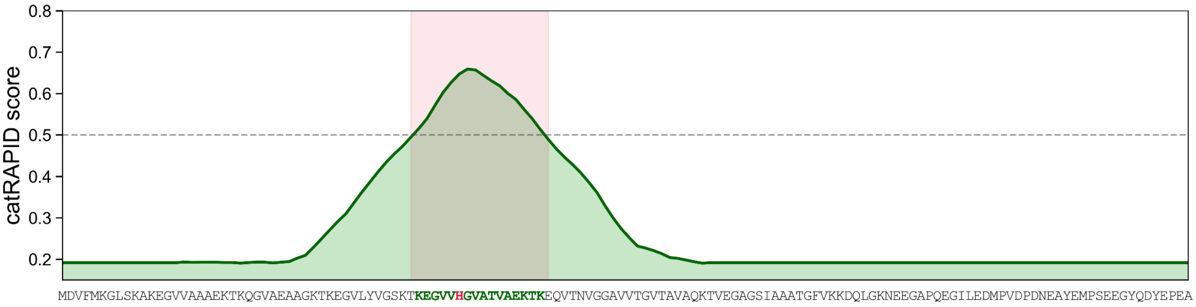


**Supplementary figure 4.** *cat*RAPID signature profile of aS with the position of His50 highlighted with the vertical line in the predicted area with the highest RNA binding propensity. The vertical line marks the z-score threshold of 0.5, indicative of significant RNA-binding propensity, with the coloured area highlighting the corresponding residues of aS. His50, the residue presumed to be involved in the fibril catalytic activity, is highlighted in red.

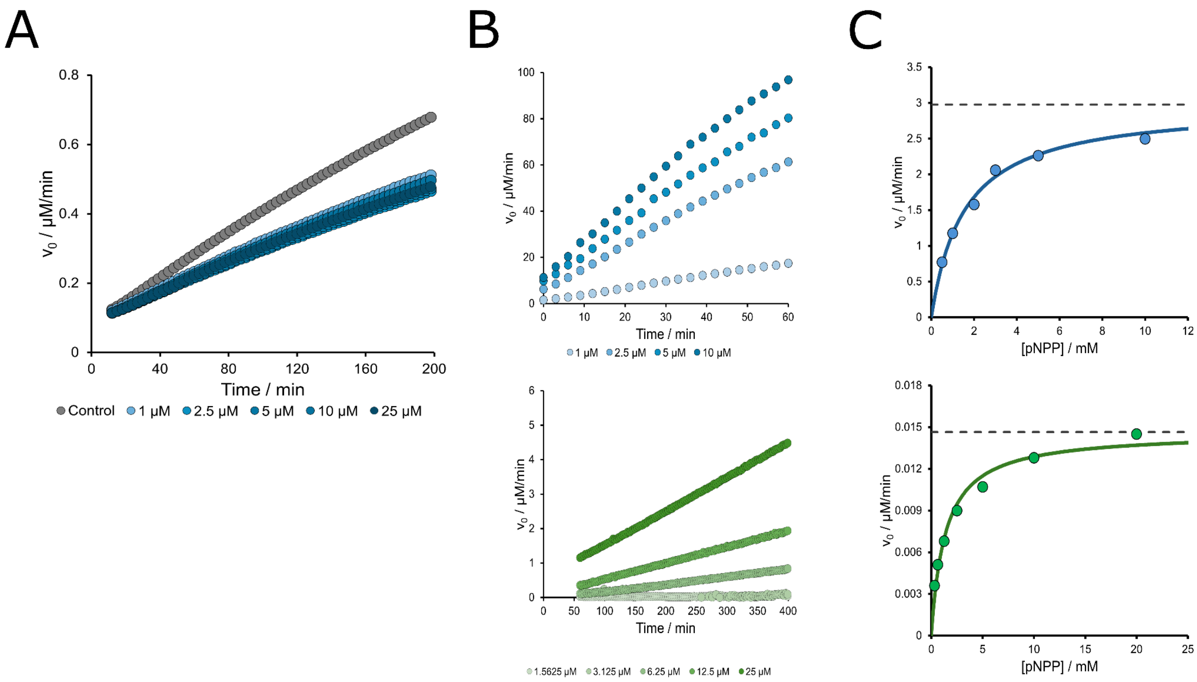


**Supplementary figure 5.** (*A*) Monomeric aS at a range of concentrations does not contribute to any pNPA cleavage compared to control (assay buffer), indicating that any observed catalytic activity is due to fibrillar aS. The rise in signal is due to water hydrolysis of pNPA, while pNPP does not hydrolyse in water and no increase in signal can be observed (data not shown). (*B*) Increasing concentrations of aS fibrils contribute to the increase in catalytic activity, further confirming the catalytic activity of aggregated aS. The initial velocities obtained from pNPA (top) and pNPP (bottom) cleavage at different substrate concentrations can be plotted to determine the catalysis parameters via the Michaelis-Menten plot (*C*). The Michaelis-Menten constant $K_M$ is determined as the concentration of the substrate at 50 % maximum reaction velocity and reflects the affinity of the enzyme to the substrate as outlined in **Materials and Methods**.

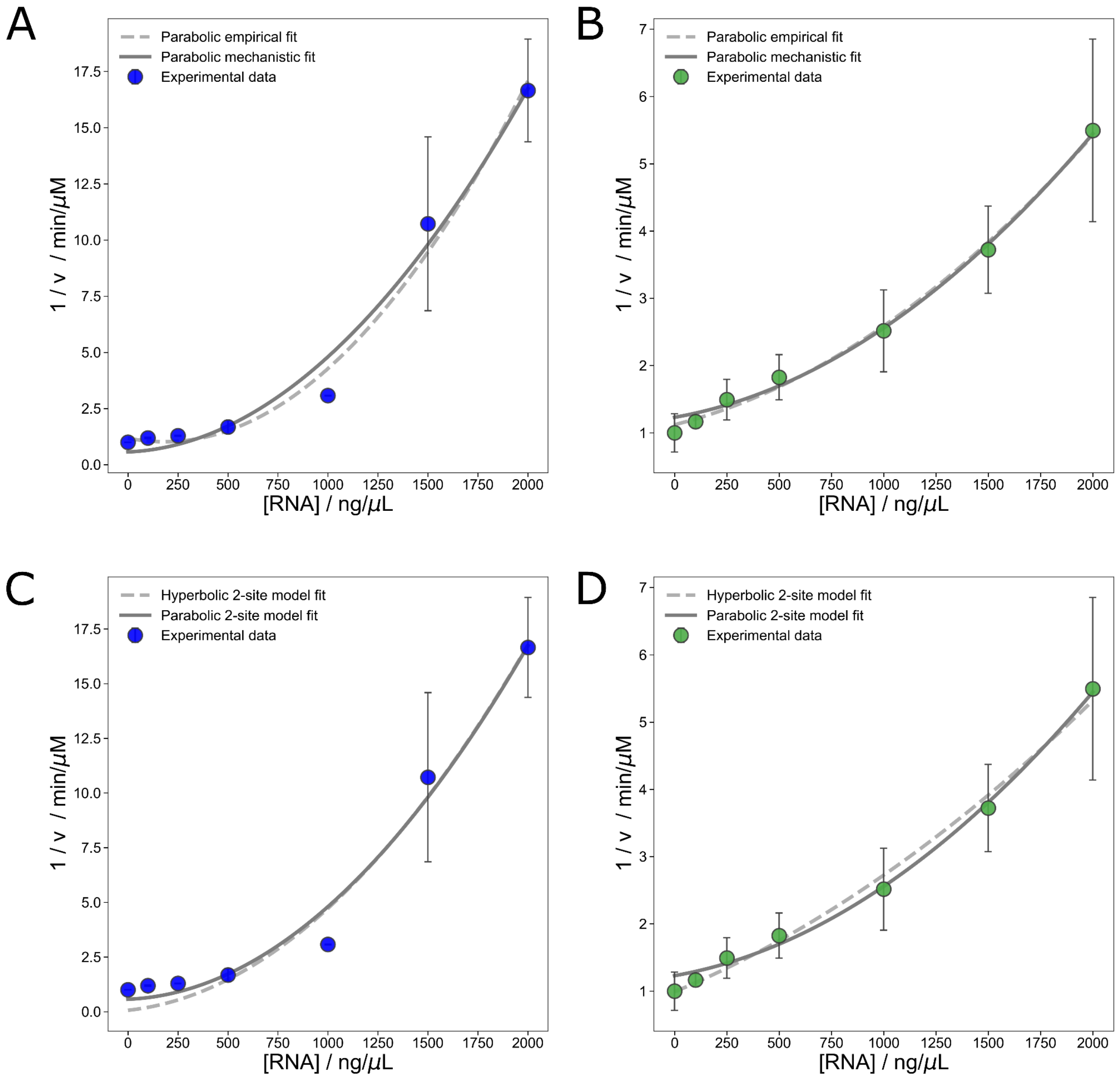


**Supplementary figure 6.** Plotting the inverse initial velocities against the inhibitor concentration such as RNA, the Dixon plot can provide insight into the mechanism of inhibition (**Supplementary Methods**). Both for pNPA (*A*) and pNPP (*B*), the exponentially increasing inverse velocities suggest either a parabolic or complex hyperbolic mechanism with multiple independent sites. The fits for both models were compared for pNPA (*C*) and pNPP (*D*) and while the parabolic two-site inhibition model produced a lower residual sum of squares (RSS) than the hyperbolic two-site model, the improvement was not statistically significant (F-test, $p > 0.05$).

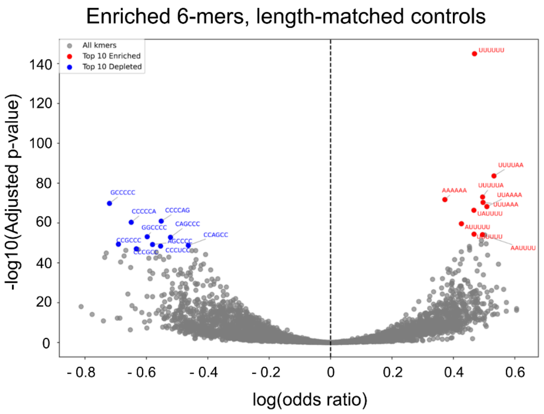


| Top 10 depleted motifs | | | | | Top 10 enriched motifs | | | | |
|---|---|---|---|---|---|---|---|---|---|
| Rank | RNA motif | Odds Ratio | p-value | Adjusted p-value | Rank | RNA motif | Odds Ratio | p-value | Adjusted p-value |
| 1 | GCCCCC | 0.486672 | 2.13E-73 | 1.46E-70 | 1 | UUUUUU | 1.598182 | 2.5E-149 | 1E-145 |
| 2 | CCCCAG | 0.575964 | 2.58E-64 | 1.17E-61 | 2 | UUUUAA | 1.702847 | 1.13E-87 | 2.31E-84 |
| 3 | CCCCCA | 0.522386 | 1.08E-63 | 4.42E-61 | 3 | UUUUUA | 1.640622 | 6.5E-77 | 8.88E-74 |
| 4 | GGCCCC | 0.550267 | 2.81E-56 | 7.67E-54 | 4 | AAAAAA | 1.451145 | 1.72E-75 | 1.76E-72 |
| 5 | CAGCCC | 0.59366 | 5.84E-56 | 1.49E-53 | 5 | UUUAAA | 1.643087 | 5.76E-74 | 4.72E-71 |
| 6 | CCGCCC | 0.501011 | 2.14E-52 | 4.61E-50 | 6 | UUAAAA | 1.663836 | 9.98E-72 | 5.84E-69 |
| 7 | AGCCCC | 0.559904 | 3.23E-52 | 6.29E-50 | 7 | UAUUUU | 1.59495 | 8E-70 | 4.1E-67 |
| 8 | CCAGCC | 0.629117 | 9.84E-52 | 1.83E-49 | 8 | AUUUUU | 1.531416 | 6.21E-63 | 2.31E-60 |
| 9 | CCCUCC | 0.574822 | 2.19E-51 | 3.91E-49 | 9 | UGUUUU | 1.595677 | 1.2E-57 | 4.1E-55 |
| 10 | CCCGCC | 0.531113 | 6.07E-50 | 9.56E-48 | 10 | AAUUUU | 1.640074 | 2.22E-57 | 6.99E-55 |

**Supplementary figure 7.** Analysis of enriched hexanucleotide motifs in protein$^{RNA}$ fibrils, matched to transcript length. Similarly to abundance-matched controls, there is a clear enrichment of AU-rich motifs and a depletion of GC-rich motifs.

### Supplementary Tables

**Supplementary table 1.** Kinetic parameters of catalytic activity of aS fibrils, compared to values reported in literature (1) . The values reported have been calculated from parameters, obtained in 7 (pNPP) and 3 independent experiments (pNPA). The fibrils were prepared freshly for each individual experiment (see Materials and Methods).

### SUPPLEMENTARY METHODS

##### Inhibition mechanism analysis and fitting

One of the most widespread methods for determining the mechanism of action for an enzymatic inhibition is through plotting the inverse reaction velocities against the concentration of the inhibition i.e. the Dixon plot. The mechanism of enzymatic inhibition is then determined according to the shape of the inhibition curve and can be fitted with the according mechanistic model of inhibition. The Dixon plot for aS fibril catalytic activity in the presence of increasing concentrations of RNA appeared to have a positive exponential curvature, indicating a complex mechanism such as parabolic inhibition (CITE). The data were then fitted both to a simpler, empiric quadratic function (Eq. 1), as well as the mechanistic parabolic inhibition equation for two independent binding sites (Eq. 2):

(Eq. 1): $F(x) = A + Bx + Cx^2$

(Eq. 2): $\frac{1}{v_i} = \frac{1}{v_{max}}\left(1 + \frac{[RNA]}{K_{i,1}} + \frac{[RNA]^2}{\left(K_{i,1} \cdot K_{i,2}\right)}\right)$

To compare the reliability of the parabolic mechanism fit compared to the hyperbolic mechanism, the data was further fitted to the mechanistic hyperbolic equation for 2 independent binding sites (Eq. 3):

(Eq. 3): $\frac{1}{v_i} = \frac{1}{v_{max}} \frac{\left(1 + \frac{[RNA]}{K_{i,1}}\right)\left(1 + \frac{[RNA]}{K_{i,2}}\right)}{\left(1 + F_{ESI}\right)}$
