## Supplementary Table 1 for "RNA interactions drive structural and functional diversification of α-Synuclein fibrils"

| | **Phosphatase (pNPP)** | | **Esterase (pNPA)** | |
|---|---|---|---|---|
| | Present work | Horvath et al. | Present work | Horvath et al. |
| **KM / mM** | 1.07 ± 0.72 | 0.5 ± 0.13 | 1.54 ± 0.44 | 4.3 ± 2.5 |
| **kcat / $s^{-1}$** | $0.9\times10^{-5} \pm 0.5\times10^{-5}$ | $30\times10^{-5} \pm 10\times10^{-5}$ | 0.005 ± 0.002 | 0.012 ± 0.003 |
| **ε / $M^{-1}$ $s^{-1}$** | 0.01 ± 0.006 | 0.6 | 3.2 ± 0.6 | 2.9 |
